# A Comprehensive Toxicological Safety Evaluation of *Anaerostipes caccae*

**DOI:** 10.1101/2025.07.08.663675

**Authors:** Vickie Modica, Róbert Glávits, Amy Clewell, John R. Endres, Gábor Hirka, Adél Vértesi, Erzsébet Béres, Ilona Pasics Szakonyiné

## Abstract

*Anaerostipes caccae* CLB101 (CLB101) is an obligate, anaerobic bacteria that was isolated from the stool of a healthy infant. Due to its ability to produce butyrate and its potential promotion of microbiome health through multiple homeostatic interactions there is interest in its consumption by humans. No toxicity data are publicly available for any strain of *A. caccae*. Therefore, its genotoxic and toxicological potential was investigated in the current study. Due to its anaerobic nature, a genotoxicity evaluation was performed using the in vivo comet assay and the in vivo mammalian micronucleus assay, which found no evidence of clastogenicity or aneugenicity. General toxicity and potential target organs were assessed in a 90-day, repeated-dose, oral toxicity study using 0, 250, 500, and 1000 mg/kg bw/day in Wistar rats. CLB101 exposure did not result in adverse effects in male or female rats when evaluated for clinical signs, body weight, food consumption, clinical pathology, organ weight, and histopathology after administration, at any dose. Therefore, a NOAEL of 1000 mg/kg bw/day, equivalent to 1.9 x 10^11^ CFU/kg bw/day, was determined in both male and female Wistar rats.

## 1. Introduction

The human microbiome contains an enormous number of microorganisms through which it contributes to health and helps defend its host against pathogens and disease by contributing to barrier function, modulating immune function and inflammation, and promoting metabolism via the break-down of non-digestible carbohydrates and the production of nutrients, conjugated bile acids, and amino acids (Riviere et al., 2016). Therefore, the discovery of safe probiotic species/strains with the potential to improve or restore the microbiome is of great interest. Of potential significance in this pursuit, the bacterium *Anaerostipes caccae* (family, *Lachnospiraceae*; genus, *Anaerostipes*) was isolated in 2000 from a healthy human fecal sample (strain L1-92)(Barcenilla et al., 2000; Schwiertz et al., 2002) and later described as non-motile, non-spore-forming, rod-shaped, obligately anaerobic, Gram-variable (generally considered Gram-positive but can stain Gram-negative in cultures older than 16 hours), catalase and oxidase-negative, and non-hemolytic (Schwiertz et al., 2002).

Perhaps its most notable and studied characteristic is its cross-feeding capability, or its ability to transform carbohydrates and the metabolites of multiple other gut microbiome species (including *Bifidobacterium* and *Lactobacillus spp.* and mucin degraders, like *Akkermansia muciniphila*), particularly lactate and acetate, into the short-chain fatty acid (SCFA) butyrate (Belzer et al., 2017; Falony et al., 2009; Schwiertz et al., 2002; Van den Abbeele et al., 2011)—a process found to drive microbial abundance in the infant gut (Chia et al., 2020; Chia et al., 2021).

*A. caccae* is a member of one of the most abundant bacterial genera found in the human microbiome (Morinaga et al., 2021) and is grouped in the Clostridial cluster XIVa within the class Clostridia, which includes anaerobic, butyrate-producing bacteria (Falony et al., 2009). Falony et al. (2009) found that when compared to other cluster XIVa butyrate producers in the gut (*Roseburia faecis*, *Roseburia hominis*, *Roseburia intestinalis*, and *Roseburia inulinivorans*), *A. caccae* was unique as the lone lactate consumer (Falony et al., 2009). As an essential metabolite and product of microbial fermentation, butyrate promotes colonic immune homeostasis by inducing the differentiation of colonic regulatory T-cells, providing energy for colonocytes, and stimulating the growth and/or activity of known health-promoting, commensal *Bifidobacteria* (Riviere et al., 2016; Tanno et al., 2019). Related to its ability to produce butyrate, several studies, in specific populations, have associated increased or decreased abundances of *A. caccae* with positive or negative outcomes, respectively (Brickman et al., 2024; Caso et al., 2021; Chia et al., 2020; Chia et al., 2021; Feehley et al., 2019; He et al., 2022).

*A. caccae* CLB101 (CLB101) is a butyrate-producing, wild-type strain that was isolated from the feces of a healthy infant (previously referred to as *A. caccae* LAHUC) (Hesser et al., 2024). Due to an interest in the use of CLB101 in food and because of the lack of safety and toxicity-related studies found in the literature, preclinical investigations to evaluate its genotoxic potential and subchronic oral toxicity in rats were performed. Because *A. caccae* is anaerobic, in vivo investigations, which can deposit the test item directly into the digestive tract, were considered more appropriate than in vitro assays, which are otherwise commonly used in toxicity batteries, but expose the test item to air (e.g., bacterial reverse mutation test, in vitro chromosomal aberration assay, and in vitro micronucleus assay). Therefore, in addition to a 90-day oral toxicity study, an in vivo comet assay was chosen to assess for mutagenicity and clastogenicity, and an in vivo mammalian micronucleus test was utilized to assess for clastogenicity and aneugenicity. The findings from this toxicological battery will provide essential data for regulatory considerations and further applications of *A. caccae* CLB101 in human health interventions.

## 2. Material and Methods

### 2.1 Test Material

The test material, a freeze-dried powder of viable *A. caccae* CLB101 (ClostraBio, Inc., Chicago, IL, USA), is composed of 1.9 x 10^10^ colony-forming units (CFU)/g at the time of batch assays. 16S rRNA evaluation, using the SILVA database, verified the identity of *A. caccae* strain CLB101 with a 99.87% alignment to its initial species assignment.

Mitsuoka buffer, made up of phosphates, Tween 80^®^, cysteine, and distilled water, and NaOH and HCl, as needed, for pH adjustment (ClostraBio, Inc., Chicago, IL, USA) was used as the vehicle in all studies. It is used as a dilution medium to preserve the maximum number of viable cells for certain anaerobic microorganisms (Champagne et al., 2011; Muto et al., 2010). Due to the obligate anaerobic nature of *A. caccae* CLB101, formulations for dosing were prepared by careful weight measurement just before each treatment and were used within two hours of preparation while being constantly stirred.

All studies were performed in compliance with their respective Organisation for Economic Co-operation and Development (OECD) guidelines for Good Laboratory Practice (GLP), except that they were performed without analytical determination on formulations (e.g., stability, homogeneity, nominal concentrations). This deviation was due to the nature of the test item and the lack of an adequate test method in the testing facility with which to comply. Instead, as stated above, the formulations were made fresh daily in Mitsuoka buffer (vehicle) using careful weight measurements just before each treatment and were used within one to two hours of preparation while being constantly stirred.

### 2.2 Care and Use of Animals

Specific pathogen-free (SPF) rats (Han:WIST; Toxi-Coop, Budapest, Hungary) were used for the in vivo comet assay (males) and 90-day repeated-dose oral toxicity study (both sexes). Male-only SPF mice (Win:NMRI;Toxi-Coop, Budapest, Hungary), one of the standard animals used for this test, internationally, were used for the in vivo mammalian micronucleus assay. The choice of these animal strains for their respective studies was due to the laboratory’s routine utilization of them in corresponding toxicological studies and, therefore, the significant accumulation of historical control data. The animal studies were approved by the Institutional Animal Care and Use Committee of Toxi-Coop Zrt. During the acclimation and treatment periods, ad libitum access was provided to food (ssniff® SM R/M-Z+H, ssniff Spezialdiäten GmbH) and potable water. The 90-day study was additionally conducted according to the National Research Council Guide for Care and Use of Laboratory Animals (National Research Council, 2011) and in compliance with the principles of the Hungarian Act 2011 CLVIII (modification of Hungarian Act 1998 XXVIII) and Government Decree 40/2013 regulating animal protection.

The Comet assay included 23 51-61-day-old healthy males weighing 267-286 g, randomized into five animals per test and vehicle control group, and three into the positive control. The animals were acclimatized for five days and housed three/cage, except the positive controls, which were housed two/cage. The 90-day study included 80 42-46-day-old healthy male and female rats, equally divided. At randomization, the males weighed 180–207 g and the females weighed 111–135 g. They were acclimatized for six days and were housed two/sex/cage. The mouse micronucleus test included 25 8-week-old healthy males. At randomization, they weighed 35.6–41.2 g, were also acclimatized for six days, and housed five/cage. Housing environmental controls included 12-hour light/dark cycles, a temperature of 22 ± 3 °C, 30–70 % humidity, and air ventilation cycles of ≥ 10 cycles/hour. All rat test and control groups included one additional contingency animal, while the high-dose mouse group included two.

### 2.3 In Vivo Comet Assay

#### 2.3.1 Good Laboratory Practice and Test Guideline, Test System, Test Material Formulation, and Concentrations

The in vivo comet assay adhered to OECD guideline 489 (OECD, 2016b) and European Commission No 2017/735 Annex Part B, B.62 (European Commission, 2017). *A. caccae* CLB101 was orally administered in doses of 500, 1000, or 2000 mg/kg bw using a treatment volume of 10 mL/kg bw twice, 24 hours apart. A preliminary, non-GLP dose range finding test determined the appropriate doses. The negative vehicle control, Mitsuoka buffer, was provided to six animals, concurrent to the test doses, and the positive control, ethyl methanesulfonate (EMS, Sigma-Aldrich) in ultrapure water (ASTM Type I) was given at a dose of 200 mg/kg bw to four animals one time on day one. Each group included one contingency animal in case a replacement animal was required during the study.

Mitsuoka buffer is rarely used in the laboratory; as such, the testing facility does not have a separate historical database for it. Therefore, initial validation and reliability studies were performed under the same conditions as the current study to determine its suitability for the target tissues. The suitability of Mitsuoka buffer was confirmed by comparison with the laboratory’s available historical control database for aqueous vehicles based on seven, ten, and nine experiments in the cases of the stomach, duodenum, and liver samples, respectively. In the current study, the percent tail DNA, tail DNA, and olive tail moments (OTM) values obtained for the target tissues were compared (by 2-Sample t-Test) to corresponding values of vehicles commonly used in the lab. As there were no significant differences, Mitsuoka buffer was considered adequate for use as a negative control.

### 2.3.2 Experimental Procedures and Evaluation of Data

Animals were weighed before each treatment and at termination. They were observed for morbidity, mortality, general symptoms, and clinical signs on day zero before the treatment started at hours one, two, and four after the initial dose, again at hours one and two after the second dose administration, and immediately before termination, 3–4 hours after the second treatment. At termination, the appearance of target tissues and organs was recorded, and cell samples were collected.

Cell suspensions were created. The glandular stomach and a sample of duodenum were separately washed, briefly incubated in mincing buffer, then scraped for cells with a scalpel and placed in the mincing buffer on ice for ∼30 seconds. A portion of the left lateral lobe of the liver was washed and then placed in mincing buffer, minced with a pair of fine scissors, and then chilled on ice for ∼30 seconds. A portion of each cell suspension was screened for cytotoxicity using the trypan blue (HyClone Laboratories Inc.) dye exclusion technique (Strober, 2001; Tice et al., 2000), which determines cell viability and provides preliminary information on the quality of cell preparation (as either genotoxicity or cytotoxicity can result in deoxyribonucleic acid (DNA) migration in an in vivo comet assay). Additionally, representative tissue samples were stored in a 4% formaldehyde (Lach:ner) solution for later histopathological examination, if needed, due to positive genotoxic in vivo comet assay results.

To prepare slides, low melting point agarose (0.5%; Sigma-Aldrich) was mixed into the individual cell preparations from which ∼10^4^–10^5^ cells were placed on slides that were pre-layered with normal melting point agarose (0.5%; Sigma-Aldrich), allowed to solidify, then covered with normal agarose and allowed to solidify again. The slides were immersed overnight in a dark refrigerator at 2–8 °C in a pH 10 lysing solution (NaCl (Lach:ner), EDTA (Merck), tris (aminomethane; Sigma-Aldrich), N-lauroylsarcosine (Sigma-Aldrich), NaOH (Acros Organics), Triton X-100 (Sigma-Aldrich) and DMSO (Sigma-Aldrich)). Subsequently, the cells were unwound (30 minutes) and then electrophoresed (30 minutes at 0.7 V/cm and 270–300 mA)—both in a controlled temperature (5 °C) electrophoresis buffer (EDTA, Merck; NaOH, Acros Organics; pH > 13), protected from sunlight. The cell remnants were neutralized (tris [hydroxymethyl] aminomethane [Sigma-Aldrich] [pH: 7.5, three times for 5 min]), then preserved (absolute ethanol [Lach:ner] for 5 min) and air-dried on slides. DNA on the preserved slides was stained with ethidium bromide (2 μg/ml, Sigma-Aldrich).

The slides were coded and blind scored—for each tissue sample, fifty cells/slide were randomly scored (150 cells/animal or 750 cells/treatment and control groups and 450/positive control). Comets were measured with a digital camera (CMOS Alpha DCM 510B) linked to a digital analyzer system using a fluorescence microscope (Nikon Upright Microscope Eclipse Ci-S) with excitation filter (TRITC). For image analysis, the Andor Komet GLP 7.1.0 (Andor Technology) was used. The independent endpoints of DNA strand breakage were measured by percent tail DNA, tail length, and OTM. Evaluation and interpretation of results were made based on percent tail DNA. Each slide was additionally examined for the presence of ghost cells (morphologically indicative of highly damaged cells)—their presence is often associated with severe genotoxicity, necrosis, and apoptosis.

The study is considered valid when the concurrent negative and positive controls, especially relative to percent tail DNA, correspond to their respective laboratory historical control ranges (within the 95% confidence intervals), the concurrent positive control produces a statistically significant increase compared with concurrent negative control, adequate numbers of cells and doses are analyzed, and the highest dose selected was according to OECD 489 guideline. The test is considered clearly negative if the test item does not induce statistically significant increases compared with concurrent negative controls, there is no dose-related increase when evaluated with an appropriate trend test, results are within the historical control range, and there is evidence of exposure to the target tissues. The test is considered clearly positive if at least one test concentration is statistically significantly increased compared with concurrent negative controls, an increase is dose-related when evaluated with an appropriate trend test, and the results are outside of the historical control range.

### 2.4 In Vivo Mammalian Micronucleus Test

#### 2.4.1 Good Laboratory Practice and Test Guideline, Test Material Formulation, Administration, and Dosing Schedule

An OECD 474 (OECD, 2016) compliant in vivo mammalian micronucleus test (MMT) was performed to evaluate the genotoxicity of *Anaerostipes caccae* CLB101 in the bone marrow of mice. The test was based on methods previously described (Salamone and Heddle, 1983), according to the laboratory’s SOPs, and performed according to GLP. A non-GLP preliminary oral toxicity test was conducted to determine the sex(es) and maximum dose for the main test.

Specific pathogen-free Win:NMRI male mice (Toxi-Coop ZRT) were used. All mice were aged eight weeks and weighed 35.6–41.2 g at the beginning of the study. Twenty-five mice were randomly divided into five groups of five, a negative and positive control, and three test groups. Two additional contingency mice were given the high dose in case of premature death(s). The test material and the negative control were prepared in Mitsuoka buffer (ClostraBio, Inc., Chicago, IL, USA). The negative control and test material were twice administered at 0, 500, 1000, or 2000 mg/kg bw by gavage in a volume of 10 mL/kg bw, 24 h apart. The positive control was administered in one intraperitoneal dose of 60 mg/kg bw of cyclophosphamide (Sigma-Aldrich) dissolved in sterile water at a volume of 10 mL/kg bw.

#### 2.4.2 Experimental Procedures and Evaluation of Data

The mice were evaluated for visible signs of reactions to treatment immediately after dosing and at regular intervals. Body weights were determined prior to dosing on each treatment day. All animals were euthanized by cervical dislocation 24 hours after their only/last treatment, and marrow samples were taken immediately from both exposed femurs. The marrow was flushed with fetal bovine serum (Sigma-Aldrich), mixed by vortex, centrifuged, and the supernatant was removed, leaving a cellular pellet that was smeared on slides. After drying at room temperature, the smears were fixed with methanol (Lach:Ner sro) for a minimum of five minutes, stained with 10% Giemsa (Merck KGaA) for 25 minutes, rinsed with distilled water, and coated with EZ-mount (Thermo Fisher). One slide from each animal was given a code number to facilitate blind scoring. Micronucleated polychromatic erythrocyte (MPCE) frequency was determined as a percentage of the first 4000 polychromatic erythrocytes counted in the optic field, and the ratio of immature to total erythrocytes was determined by counting a total of at least 500 erythrocytes as a measure of potential toxicity and target cell exposure. The results were evaluated per the OECD 474 guideline.

### 2.5 90-Day Repeated-Dose Oral Toxicity Study in Rats

#### 2.5.1 Good Laboratory Practice and Test Guideline, Test Material Formulation, Administration, and Dosing Schedule

A GLP, 90-day repeated-dose oral toxicity study in rats was conducted in general accordance with OECD 408 (OECD, 2018) and in accordance with the laboratory’s SOPs. Eighty healthy, specific pathogen-free Wistar rats (Toxi-Coop ZRT) were used in the experiment. Groups of two male and two female animals were housed together and acclimatized for six days in a controlled environment (22 ± 3 °C, 30–70% relative humidity, 12-hour light/dark (except on ophthalmological exam days)). The rats were divided into four groups of 10 males and 10 females each at random, with equal weight distributions among the groups. Doses of 550, 1100, and 2200 mg/kg bw/day of the test material were administered by gavage on Days 0–4 then the test doses were changed to 250, 500, and 1000 mg/kg bw/day for the duration of the study using a constant dosing volume of 10 mL/kg bw. The initial doses were chosen based on the doses used in a preceding 14-day repeated-dose oral toxicity study (unpublished), then, at the request of the sponsor and based on the expected human dosing regimen, were lowered from Day five on. Daily doses were given to eighty 42–46-day-old male and female animals with starting weights of 180–207 and 111–135 g, respectively.

#### 2.5.2 Observations, Measurements, and Examinations

Body weights were measured twice and clinical observations collected three times during the acclimatization period and on day zero. Animals were weighed twice/week during weeks 1–4, once/week for the duration of the study, and immediately before the animals were euthanized. Weekly food consumption was measured, and weekly food efficiency was calculated. Cage-side clinical observations were made during and after each treatment. More detailed hands-on clinical observations were documented the day before treatment started and then once/week. Mortality and morbidity checks were done twice each day. A functional observational battery (FOB; a modification of the procedure described by Irwin (Irwin, 1968)) was completed on Day 85. All animals received ophthalmological exams before the first treatment, and a follow-up was done for those in the control and high-dose groups on Day 85. On the day of necropsy, Day 91, vaginal smears were taken and stained with 1 % aqueous methylene blue solution (Acros Organics).

The animals were fasted for approximately 16 hours after their last treatment and were placed under anesthesia using Isofluran CP^®^ (Medicus Partner Kft.), and blood samples were taken from the retro-orbital venous plexus for clinical pathology (Days 90 for males and 91 for females). Immediately after, the anesthetized animals were sacrificed by exsanguination from the abdominal aorta. Three blood samples were taken for hematologic and clinical chemistry parameters. At necropsy, gross pathological examinations were conducted on each animal, including organ weight measurements and removals, and tissues were preserved in a 4 % formaldehyde solution (Lach-Ner sro), except the testes and epididymides which were fixed for one day in a modified Davidson solution (prepared in the laboratory) and then preserved in the formaldehyde solution for future examination, if needed. The preserved and fixed tissues were trimmed, processed, embedded in paraffin (DiaPath S.p.A.), sectioned with a microtome (DiaPath S.p.A.), placed on standard microscope slides, stained with hematoxylin (DiaPath S.p.A.) and eosin (DiaPath S.p.A.), and examined by light microscopy. All control and high-dose group animals received full histological exams on their preserved organs as well as analyses on the male reproductive organs, including organ morphology and sperm enumeration, morphology, and motility. Additionally, low and mid-dose animals with gross findings were also examined histopathologically.

### 2.6 Statistics

Statistical analyses were performed with SPSS PC+ software (SPSS, Inc., version 4), and Microsoft Excel (Microsoft) was used to check for linear trends. A P-value of < 0.05 was considered statistically significant in all tests.

#### 2.6.1 In Vivo Comet Assay

Levene’s test for equality of variances was used to assess the homogeneity of variance between groups, and the Kolmogorov–Smirnov test was utilized to assess for normal distribution of data. Based on their results, the inter-group comparisons between the vehicle control value and the corresponding values of treatment groups were performed using non-parametric Mann-Whitney U test.

#### 2.6.2 In Vivo Mammalian Micronucleus Test

The statistical frequency of observed MPCEs in the test and positive control groups were compared to the concurrent negative control group and laboratory historical control levels. A linear trend for the frequency of MPCEs was tested using adequate regression analysis with Microsoft Excel.

#### 2.6.3 90-Day Repeated-Dose Oral Toxicity Study in Rats

Analyses of body weight, food consumption, clinical pathology parameters, and organ weights were done. The heterogeneity of variance between groups was checked using Bartlett’s homogeneity of variance test. If statistically significant variance was found, the normal distribution of data was followed by the Kolmogorov-Smirnov test that, when a non-normal distribution was found, was itself followed up with non-parametric Kruskal-Wallis ANOVA. If the non-parametric Kruskal-Wallis ANOVA was positive, the Mann-Whitney U-test was performed for inter-group comparisons. If Bartlett’s found no statistically significant variance, it was followed up with a one-way ANOVA, which, when positive, was reconciled with Duncan’s Multiple Range test to assess the inter-group differences for significance. For the comparison of data in two groups, the homogeneity of variance between groups was checked by F-test, and, depending on the result, pooled or separate variance estimates of the two-sample t-test were performed. Frequencies of toxic response, ophthalmoscopy, gross pathological, and histopathological findings by sex and dose were calculated. Statistical analyses were completed separately for male and female animals.

## 3. Results

### 3.1 In Vivo Comet Assay

No mortality, clinical signs, or other toxicological signs were observed in any group after treatment with the test item or the controls. No gross pathological abnormalities, signs of tissue toxicity, or local test item effects were observed upon sample collection in any group. Throughout the study, body weight gains were within the expected parameters for normal development based on the historical control ranges for all treatment and control groups.

The percent tail DNA for all target tissues, obtained with Mitsuoka buffer as the negative (vehicle) control, were comparable to corresponding values of usual vehicles using the two-sample t-test. The duodenum and liver percent tail DNA values of the concurrent negative controls were within the 95% confidence intervals of the historical negative control data for aqueous vehicles. The stomach values were below this range, but when examined in the context of the test facility’s more robust historical control database, built for sunflower oil, the lower levels fell within the 95% confidence interval and, thus, were deemed acceptable. The range of percent tail DNA values for all tissues fell within the acceptable 1–20% proposed by the Japanese Center for the Validation of Alternative Methods validation trial (OECD, 2014). Further, the liver values were aligned with OECD 489 laboratory proficiency recommendations and published literature (Pant et al., 2014; Vasquez, 2010). Relative to the negative control, there were no statistically significant changes in the percent tail DNA in any treatment group or any tissue type. Analysis of tail lengths and OTMs of the negative control for the three tissue types mirrored the conclusion above, except that all stomach (group mean and individual) and three individual liver OTM values were also below the 95% confidence intervals of the historical negative control data for aqueous vehicles. As done for the percent tail DNA stomach values, comparisons to the sunflower oil historical control database found these lower values acceptable as they fell within the 95% confidence intervals. Regarding the concurrent positive control, all of the group mean values for percent tail DNA, the tail length, and OTM were comparable to the historical control database and were statistically significantly increased in all target tissues relative to the negative control. Two individual liver means fell above the 95% confidence limit (one each percent tail DNA and tail length); both were considered favorable as they indicated a higher EMS effect. Therefore, this in vitro comet assay was considered valid—it fulfilled all validity criteria regarding the concurrent negative and positive controls and, as discussed previously, the number of cells analyzed, and the high-dose test group were based on OECD 489 guidelines. The percentages of ghost cells in all dose group tissue samples were within the interval of the laboratory’s historical control data, and all of the test group mean values were within the historical control range. Individual means for the liver samples were also within the historical control range. Some stomach and duodenum individual mean values were slightly lower, which was considered favorable, highly acceptable (as they indicated good cell suspension quality without test item effect), and without effect on the study results. Cytotoxicity screening using the Trypan blue exclusion method resulted in no difference in the frequency or severity of coloration between the treated cells and the control cells. A histopathological assessment was deemed unnecessary due to the lack of genotoxicity in all tissue samples. See Table 1 for the in vivo comet assay test results.

**Table 1.** Summary of *A. caccae* CLB101 Relevant Comet Assay Findings in Stomach, Duodenum, and Liver.

| Test | 0 mg/kg<br>bw/day | 500 mg/kg<br>bw/day | 1000 mg/kg<br>bw/day | 2000 mg/kg<br>bw/day | EMS<sup>†</sup> |
| --- | --- | --- | --- | --- | --- |
| **Stomach** |  |  |  |  |  |
| Mean Median % Tail DNA<sup>‡</sup> | 3.18<sup>a</sup> | 3.94<sup>a</sup> | 4.35<sup>a</sup> | 4.16<sup>a</sup> | 25.75 |
| SD | 0.47 | 1.00 | 1.1 | 1.03 | 3.59 |
| SS |  | NS | NS | NS | * |
| historical control range | 4.52–13.09 |  |  |  | 14.34–37.19 |
| Tail Length (µm) | 7.53 | 7.73 | 8.07 | 8.62 | 50.43 |
| SD | 0.34 | 0.40 | 1.57 | 1.97 | 1.31 |
| SS |  | NS | NS | NS | * |
| Historical control range | 6.17–56.77 |  |  |  | 28.55–75.56 |
| Olive Tail Moment | 0.34<sup>b</sup> | 0.38<sup>b</sup> | 0.43 | 0.42 | 5.50 |
| SD | 0.04 | 0.05 | 0.1 | 0.09 | 0.92 |
| SS |  | NS | NS | NS | * |
| historical control range | 0.42–4.68 |  |  |  | 1.91–10.16 |
| Percent ghost cells | 4.17 | 4.37 | 4.53 | 4.26 | 11.52 |
| SD | 0.95 | 1.99 | 1.44 | 2.03 | 0.35 |
| SS |  | NS | NS | NS | * |
| historical control range | 4–14% |  |  |  | 12–25% |
| relative ratio of ghost cells:control |  | 1.05 | 1.09 | 1.02 | 2.76 |
| **Duodenum** |  |  |  |  |  |
| Mean Median % Tail DNA<sup>‡</sup> | 3.74 | 3.73 | 5.01 | 3.39 | 25.33 |
| SD | 0.94 | 0.27 | 1.56 | 0.64 | 3.83 |
| SS |  | NS | NS | NS | * |
| historical control range | 0.47–16.88 |  |  |  | 10.04–36.09 |
| Tail Length (µm) | 7.34 | 7.15 | 8.69 | 8.47 | 54.25 |
| SD | 0.34 | 0.70 | 2.20 | 2.38 | 8.02 |
| SS |  | NS | NS | NS | * |
| historical control range | 0.00–56.67 |  |  |  | 17.66–78.74 |
| Olive Tail Moment | 0.35 | 0.36 | 0.46 | 0.34 | 5.13 |
| SD | 0.06 | 0.04 | 0.14 | 0.07 | 1.15 |
| SS |  | NS | NS | NS | * |
| historical control range | 0.00–5.08 |  |  |  | 0.04–10.15 |
| Percent ghost cells | 3.06 | 2.93 | 4.34 | 4.01 | 3.83<sup>c</sup> |
| SD | 1.6 | 1.53 | 2.86 | 2.71 | 1.07 |
| SS |  | NS | NS | NS | NS |
| historical control range | 3–11% |  |  |  | 4–24% |
| relative ratio of ghost cells:control |  | 0.96 | 1.42 | 1.31 | 1.25 |
| **Liver** |  |  |  |  |  |
| Mean Median % Tail DNA<sup>‡</sup> | 3.55 | 3.3 | 3.41 | 3.79 | 22.04 |
| SD | 1.11 | 1.00 | 0.65 | 1.34 | 4.63 |
| SS |  | NS | NS | NS | * |
| historical control range | 2.82–7.93 |  |  |  | 10.03–27.00 |
| Tail Length (µm) | 7.78 | 7.13 | 7.17 | 7.22 | 53.08 |
| SD | 0.81 | 0.56 | 0.73 | 0.67 | 11.07 |
| SS |  | NS | NS | NS | * |
| historical control range | 1.11–40.60 |  |  |  | 27.97–63.63 |
| Olive Tail Moment | 0.4 | 0.34<sup>d</sup> | 0.33<sup>d</sup> | 0.36<sup>d</sup> | 4.39 |
| SD | 0.12 | 0.07 | 0.06 | 0.11 | 1.65 |
| SS |  | NS | NS | NS | * |
| historical control range | 0.40–2.43 |  |  |  | 1.11–6.87 |
| Percent ghost cells | 1.04 | 1.16 | 1.29 | 1.04 | 2.58 |
| SD | 0.57 | 0.82 | 1.1 | 0.35 | 0.62 |
| SS |  | NS | NS | NS | * |
| historical control range | 1–6% |  |  |  | 1–10% |
| relative ratio of ghost cells:control |  | 1.12 | 1.24 | 1.00 | 2.48 |

<sup>‡</sup>performed in 750 cells (450 cells in EMS); Historical control ranges reported as 95% confidence intervals.
\*statistically significant, p=0.05, Mann-Whitney U-test versus control.
Abbreviations: DNA, deoxyribonucleic acid; EMS, Ethyl methanesulfonate; NS, not significant; SD, standard
<sup>†</sup>200 mg/kg (positive control)
<sup>a</sup>The lower % Tail DNA stomach values were also examined in context of more robust databases of another vehicle (sunflower oil). The group mean as well as the individual means fell well within the 95% confidence intervals, C-charts (1.15–14.79).
<sup>b</sup>The lower % Olive Tail Moment stomach values were examined in context of more robust databases of another vehicle (sunflower oil). The group mean as well as the individual means fell well within the 95% confidence intervals, C-charts (0–4.12).
<sup>c</sup>Group mean result was rounded to 4% for the interpretation. Note that there was one individual lower mean of 3 % (below the 4–24 % historical control range), but that EMS does not typically increase ghost cell frequency, therefore high frequencies are not expected.
<sup>d</sup>The lower % Olive Tail Moment liver values were examined in context of more robust databases of another vehicle (sunflower oil). The group mean as well as the individual means fell well within the 95% confidence intervals, C-charts (0.19–1.87).

### 3.2 In Vivo Mammalian Micronucleus Test

In a preliminary toxicity test, no mortality, adverse reactions, or sex differences were observed in male and female Win:NMRI mice given two 2000 mg/kg bw doses of the test material. Similarly, in the main study, no animals died, and no adverse reactions were observed. The concurrent negative and positive controls produced results within their respective historical control ranges, with a large statistically significant increase in the number of MPCEs in the positive control group mice relative to the negative; thus, the study was considered valid.

No statistically significant increases in MPCEs were observed in any treatment group compared to the concurrent negative control group, and the MPCEs that were found were compatible with the negative historical control data. A statistically lower proportion of immature to total erythrocytes (-11%) was observed, demonstrating exposure of the test item to the bone marrow. See Table 2 for the in vivo mammalian micronucleus test results.

**Table 2.** Results of *A. caccae* CLB101 In vivo Mammalian Micronucleus Test.

| Groups (n =5<sup>†</sup>) | Sampling time<br>(hours<br>following final<br>treatment) | Total number<br>of PCEs<br>analysed | MPCE<br>(per 4000 PCE) |  | PCE/<br>PCE+NCE |  |
| --- | --- | --- | --- | --- | --- | --- |
|  |  |  | mean | SD | mean | SD |
| Concurrent Negative<br>(Vehicle) Control | 24 | 20,000 | 6.20 | 1.64 | 0.54 | 0.03 |
| Historical Negative<br>Control (95% control<br>interval) | 24 | 880,000 | 3.32–7.59 | — | 0.506–0.547 | — |
| 500 mg/kg bw | 24 | 20,000 | 6.60 | 1.67 | 0.50*U | 0.03 |
| 1000 mg/kg bw | 24 | 20,000 | 6.00 | 1.58 | 0.50*U | 0.02 |
| 2000 mg/kg bw | 24 | 20,000 | 6.60 | 1.67 | 0.48*DN,**U | 0.01 |
| Positive Control<br>(60mg/kg bw) | 24 | 20,000 | 150.80**U | 10.92 | 0.42**DN, **U | 0.05 |

Abbreviations: MPCE, micronucleated polychromatic erythrocytes; NCE, normochromatic erythrocytes; PCE, polychromatic erythrocytes.
†Historical Negative Control (n = 44)
PCE+NCE=500
Negative (Vehicle) Control = Mitsuoka buffer
Positive Control = 60 mg/kg bw Cyclophosphamide
U: Mann-Whitney U-test Versus Control
DN: Duncan's multiple range test
The regression analysis did not show significance compared to the vehicle control
MPCE: \*\* = p < 0.01 (U) to the vehicle and historical control
PCE/PCE+NCE:\*/\*\* = p < 0.05/ p < 0.01 (DN) to the vehicle control
PCE/PCE+NCE:\*/\*\* = p < 0.05/ p < 0.01 (U/) to the historical control

### 3.3 90-Day Repeated-Dose Oral Toxicity Study in Rats

#### 3.3.1 Mortality, Clinical Observations, and Ophthalmology

No mortality occurred in any group. There were no adverse clinical signs or unusual behavior observed during the daily or detailed, weekly clinical examinations. No abnormal behavior or reactions to different types of stimuli or changes in physical state were observed in the control or high-dose animals relative to the control on Day 85 during the FOB. Additionally, there were no changes to the normal eye findings from just before the start until the end of the treatment period in any male or female animal in the high-dose or control groups.

#### 3.3.2 Body Weights and Food Consumption

There were no statistically significant differences in the mean body weights or body weight gain, or mean daily food consumption in either sex, relative to the concurrent negative control. Most test groups experienced transient weight gain fluctuations; despite this, the mean body weights were similar for the entire treatment period, and there were no statistically significant changes to the mean body weights or overall body weight development, as shown in Figure 1.

**Figure 1.**
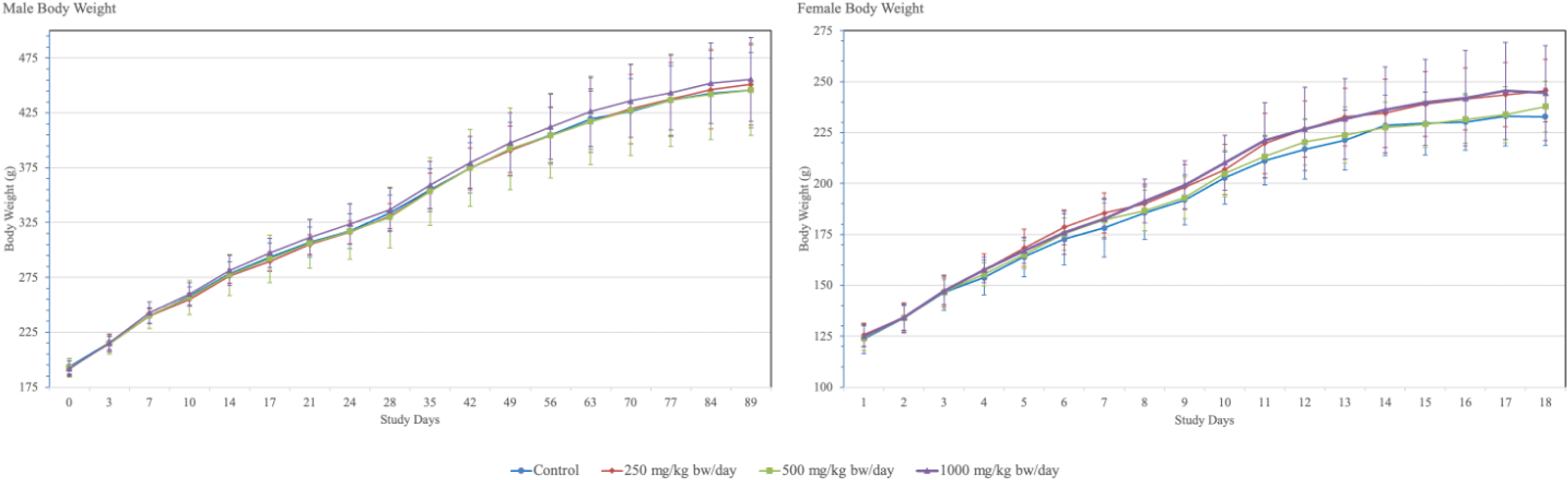
Male and Female Body Weight Development. (a) Male body weights (b) Female body weights.

The mean daily food consumption was not adversely affected in any group during the 90- day treatment period. However, the mid-dose group females consumed statistically significantly more on days 63–70, and all female test groups consumed statistically significantly more on days 70–77, while the mid-dose group consumed more on days 63–70. Despite the increased food consumption in these groups, their body weight gain was not statistically significantly affected during the same weeks. See supporting information, Tables S1 and S2 for food consumption and body weights, respectively.

#### 3.3.3 Clinical Pathology

No adverse changes were seen in the clinical pathology analyses overall, but there were some minor, yet statistically significant differences, relative to the control. These include various hematology parameter findings in male (high-dose) and female (all) test groups, and various clinical chemistry findings, including thyroid-related findings, in all male and female test groups. See the hematology and clinical chemistry with thyroid Tables 3 & 4, respectively, and section 4 for a relevant discussion. Note that one low-dose female animal was mistakenly over-anesthetized before blood sampling; thus, it was not included in the clinical pathology results.

**Table 3.**
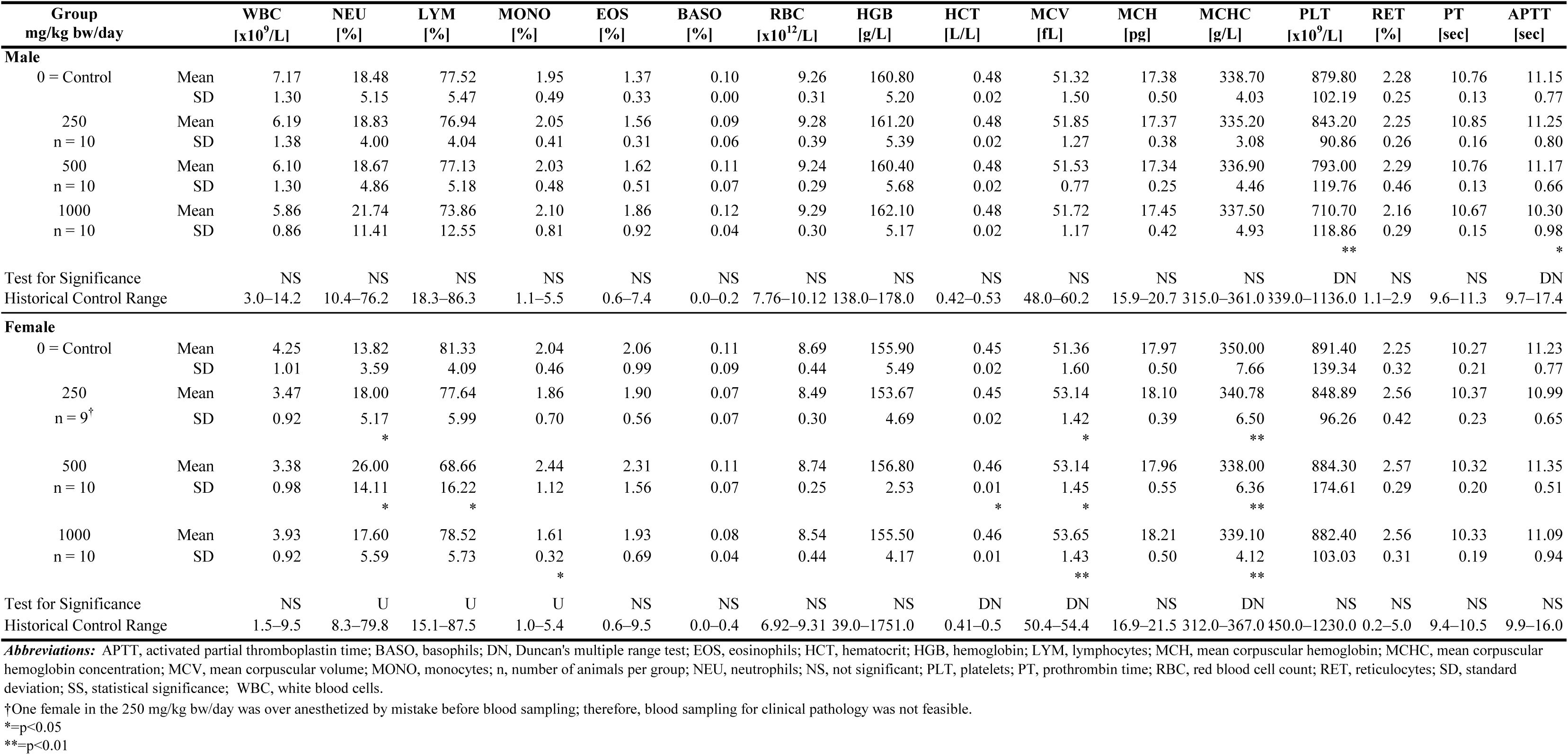
*A. caccae* CLB101 90-day Hematology.

**Table 4.**
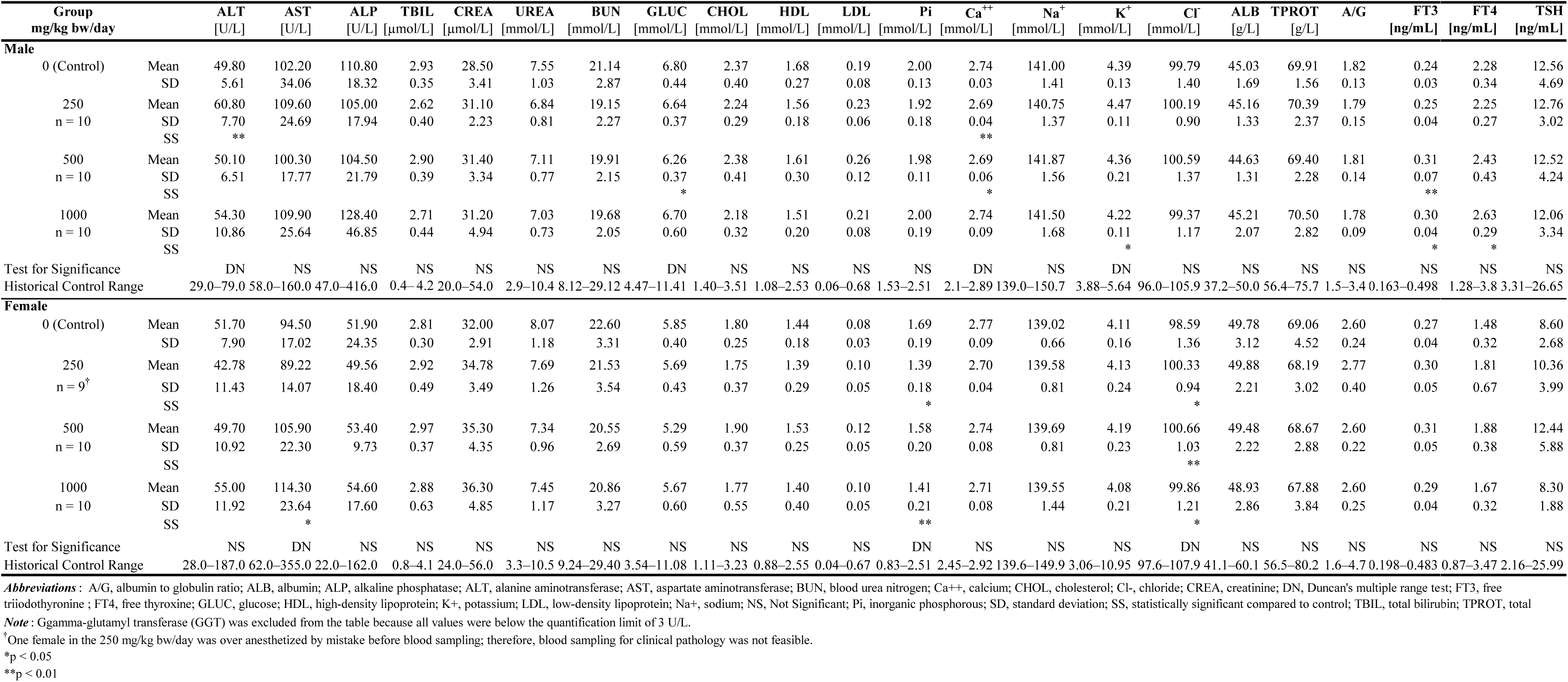
*A. caccae* CLB101 90-day Clinical Chemistry with Thyroid.

#### 3.3.4 Necropsy, Organ Weights, and Histopathology

There were no test item-related necropsy findings in any test group relative to the control. However, there were non-specific findings in all control and test groups (see Table 5). Pyelectasia predominated with incidence in all groups. Additionally, one high-dose male had brownish-red colored lymph nodes, all female groups had some level of hydrometra, two mid-dose females had a reddish-brown thymus, the high-dose groups had one animal with larger than normal adrenal glands and two with smaller than normal and pale ovaries, while in the control group hernia diaphragmatica (with liver that spread into it) was also found. The weights of all the examined organs were comparable to those of the controls for all male and female test groups. There were some minor findings in all the female test groups, including statistically significant, dose-dependent decreases in the weights of thymus glands (see Table 6 for organ weights and Tables 7 and 8 for organ weights relative to body and brain weights, respectively). Histopathological exams of the control and high-dose animals did not find test-item related lesions in the tissues or organs; those found upon gross examination were determined to have no correlative degeneration, inflammation, or fibrosis (see Table 9).

**Table 5.** *Anaerostipes caccae* CLB101 90-day Necropsy Summary.

| Organs | Observations | 0 (Control)<br>mg/kg bw/day | 250<br>mg/kg bw/day | 500<br>mg/kg bw/day | 1000<br>mg/kg bw/day |
| --- | --- | --- | --- | --- | --- |
| **Male** |  |  |  |  |  |
|  | No macroscopic findings | 7/10 | 8/10 | 9/10 | 8/10 |
| Kidneys | Dilation of the renal pelvis | 3/10 | 2/10 | 1/10 | 2/10 |
| Submandibular lymph nodes | Brownish-red colored | 0/10 | 0/10 | 0/10 | 1/10 |
| **Female** |  |  |  |  |  |
|  | No macroscopic findings | 5/10 | 3/10 | 3/10 | 4/10 |
| Adrenal glands | Larger than normal | 0/10 | 0/10 | 0/10 | 1/10 |
| Uterus | Dilation of the uterine lumen | 2/10 | 6/10 | 5/10 | 3/10 |
| Kidneys | Dilation of the renal pelvis | 2/10 | 1/10 | 2/10 | 1/10 |
| Diaphragm | Hernia 1x0,8 cm | 1/10 | 0/10 | 0/10 | 0/10 |
| Liver | Spread into the diaphragmatic hernia | 1/10 | 0/10 | 0/10 | 0/10 |
| Thymus | Brownish reddish colored | 0/10 | 0/10 | 2/10 | 0/10 |
| Ovaries | Smaller than normal, pale | 0/10 | 0/10 | 0/10 | 2/10 |

*Note* : Data represent the number of animals with observation / number of animals examined.

**Table 6.**
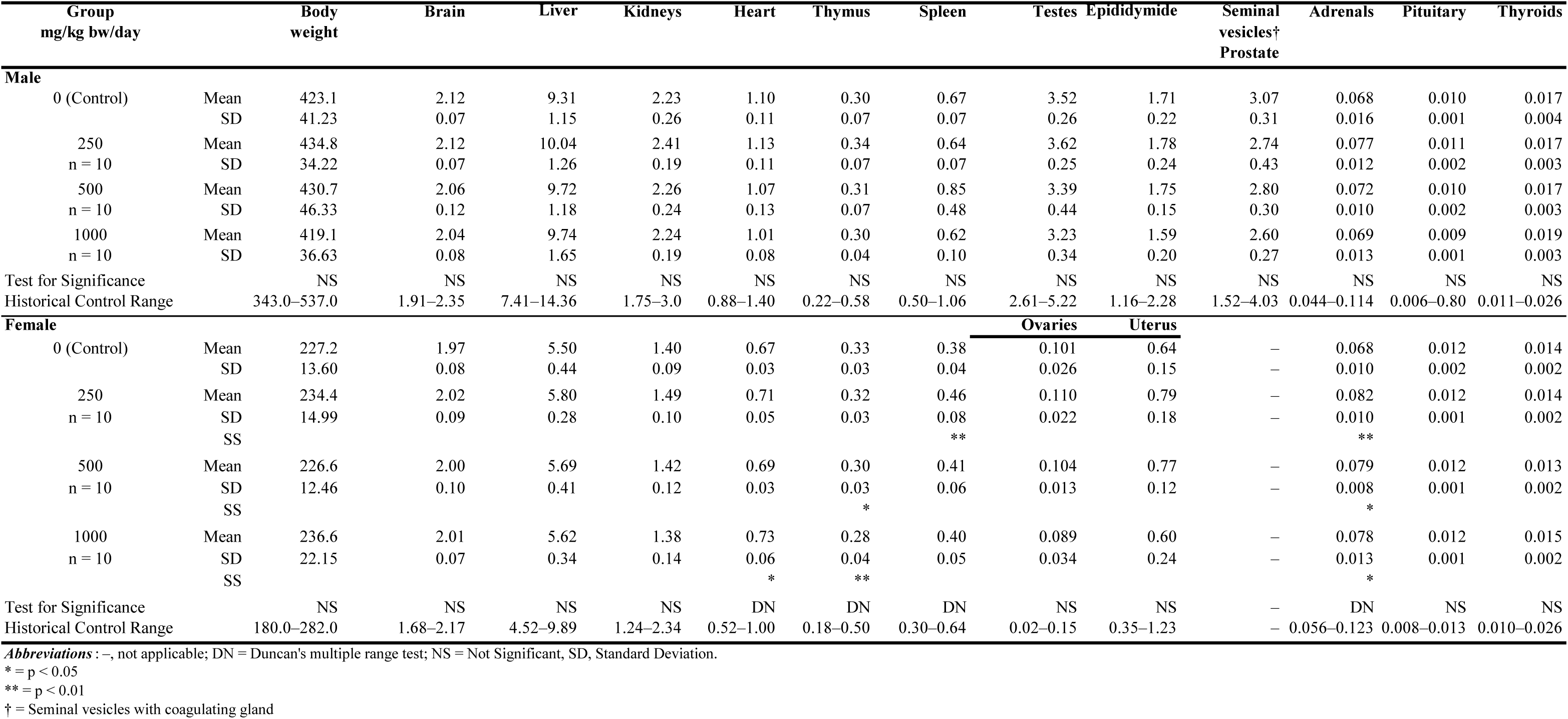
Anaerostipes caccae CLB101 90-day Organ Weights Summary.

**Table 7.** *Anaerostipes caccae* CLB101 Organ Weight Relative to Body Weight.

| Group<br>mg/kg bw/day |  | Brain | Liver | Kidneys | Heart | Thymus | Spleen | Testes | Epididymid | Seminal<br>vesicles†<br>Prostate | Adrenals | Pituitary | Thyroids |
| --- | --- | --- | --- | --- | --- | --- | --- | --- | --- | --- | --- | --- | --- |
| **Male** |  |  |  |  |  |  |  |  |  |  |  |  |  |
| 0 (Control) | Mean | 0.504 | 2.386 | 0.537 | 0.253 | 0.081 | 0.161 | 0.813 | 0.364 | 0.637 | 0.0163 | 0.0022 | 0.0037 |
|  | SD | 0.039 | 0.185 | 0.040 | 0.027 | 0.016 | 0.015 | 0.065 | 0.040 | 0.075 | 0.0022 | 0.0003 | 0.0006 |
| 250<br>n = 10 | Mean | 0.508 | 2.260 | 0.524 | 0.254 | 0.089 | 0.150 | 0.833 | 0.380 | 0.683 | 0.0148 | 0.0023 | 0.0037 |
|  | SD | 0.038 | 0.153 | 0.032 | 0.020 | 0.015 | 0.014 | 0.083 | 0.043 | 0.099 | 0.0024 | 0.0002 | 0.0003 |
| 500<br>n = 10 | Mean | 0.503 | 2.293 | 0.543 | 0.252 | 0.073 | 0.153 | 0.832 | 0.388 | 0.651 | 0.0155 | 0.0022 | 0.0038 |
|  | SD | 0.046 | 0.175 | 0.032 | 0.023 | 0.009 | 0.014 | 0.067 | 0.044 | 0.092 | 0.0015 | 0.0002 | 0.0005 |
| 1000<br>n = 10 | Mean | 0.502 | 2.292 | 0.538 | 0.261 | 0.076 | 0.161 | 0.852 | 0.374 | 0.622 | 0.0162 | 0.0022 | 0.0040 |
|  | SD | 0.041 | 0.176 | 0.031 | 0.018 | 0.011 | 0.021 | 0.075 | 0.043 | 0.079 | 0.0031 | 0.0002 | 0.0005 |
| Test for Significance |  | NS | NS | NS | NS | NS | NS | NS | NS | NS | NS | NS | NS |
| Historical Control Range |  | 0.376–0.620 | 1.998–3.156 | 0.428–0.672 | 0.206–0.349 | 0.053–0.129 | 0.109–0.241 | 0.621–1.097 | 0.270–0.518 | 0.397–0.966 | 0.011–0.025 | 0.002–0.018 | 0.002–0.007 |
| **Female** |  |  |  |  |  |  |  |  |  |  |  |  |  |
| 0 (Control) | Mean | 0.870 | 2.424 | 0.615 | 0.295 | 0.145 | 0.168 | 0.0445 | 0.283 | – | 0.0298 | 0.0051 | 0.0063 |
|  | SD | 0.081 | 0.174 | 0.042 | 0.015 | 0.012 | 0.019 | 0.0116 | 0.064 | – | 0.0040 | 0.0005 | 0.0009 |
| 250<br>n = 10 | Mean | 0.866 | 2.483 | 0.636 | 0.305 | 0.134 | 0.196 | 0.0472 | 0.337 | – | 0.0353 | 0.0053 | 0.0061 |
|  | SD | 0.063 | 0.190 | 0.064 | 0.022 | 0.012 | 0.038 | 0.0102 | 0.081 | – | 0.0051 | 0.0005 | 0.0007 |
| SS |  |  |  |  |  |  |  |  |  |  | * |  |  |
| 500<br>n = 10 | Mean | 0.882 | 2.513 | 0.631 | 0.303 | 0.131 | 0.182 | 0.0461 | 0.341 | – | 0.0351 | 0.0054 | 0.0058 |
|  | SD | 0.036 | 0.159 | 0.068 | 0.020 | 0.016 | 0.034 | 0.0070 | 0.060 | – | 0.0044 | 0.0007 | 0.0007 |
| SS |  |  |  |  |  | * |  |  |  |  | * |  |  |
| 1000<br>n = 10 | Mean | 0.853 | 2.384 | 0.583 | 0.310 | 0.119 | 0.169 | 0.0385 | 0.256 | – | 0.0330 | 0.0051 | 0.0063 |
|  | SD | 0.072 | 0.160 | 0.050 | 0.035 | 0.015 | 0.015 | 0.0150 | 0.108 | – | 0.0041 | 0.0007 | 0.0010 |
| SS |  |  |  |  |  | ** |  |  |  | – |  |  |  |
| Test for Significance |  | NS | NS | NS | NS | NS | NS | NS | NS | – | NS | DN | NS |
| Historical Control Range |  | 0.692–1.022 | 2.127–4.173 | 0.525–0.951 | 0.232–0.422 | 0.078–0.208 | 0.139–0.279 | 0.008–0.070 | 0.148–0.532 | – | 0.022–0.050 | 0.003–0.021 | 0.004–0.012 |

**Abbreviations:** –, not applicable; DN = Duncan's multiple range test; NS = Not Significant; SD, Standard Deviation.
\* = $p < 0.05$
† = Seminal vesicles with coagulating gland;

**Table 8.**
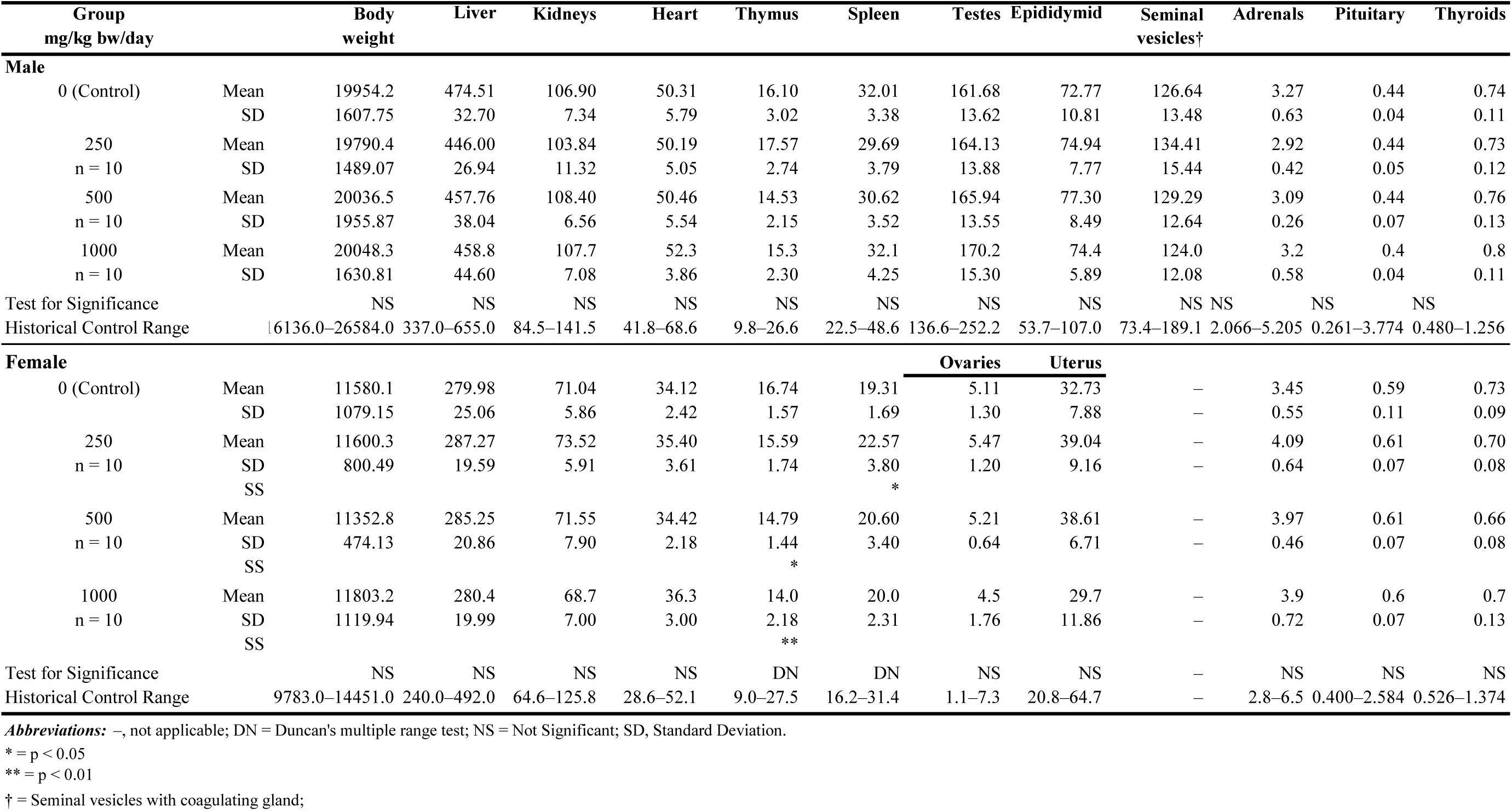
*Anaerostipes caccae* CLB101 Organ Weights and Body Weights Relative to Brain Weights.

**Table 9.**
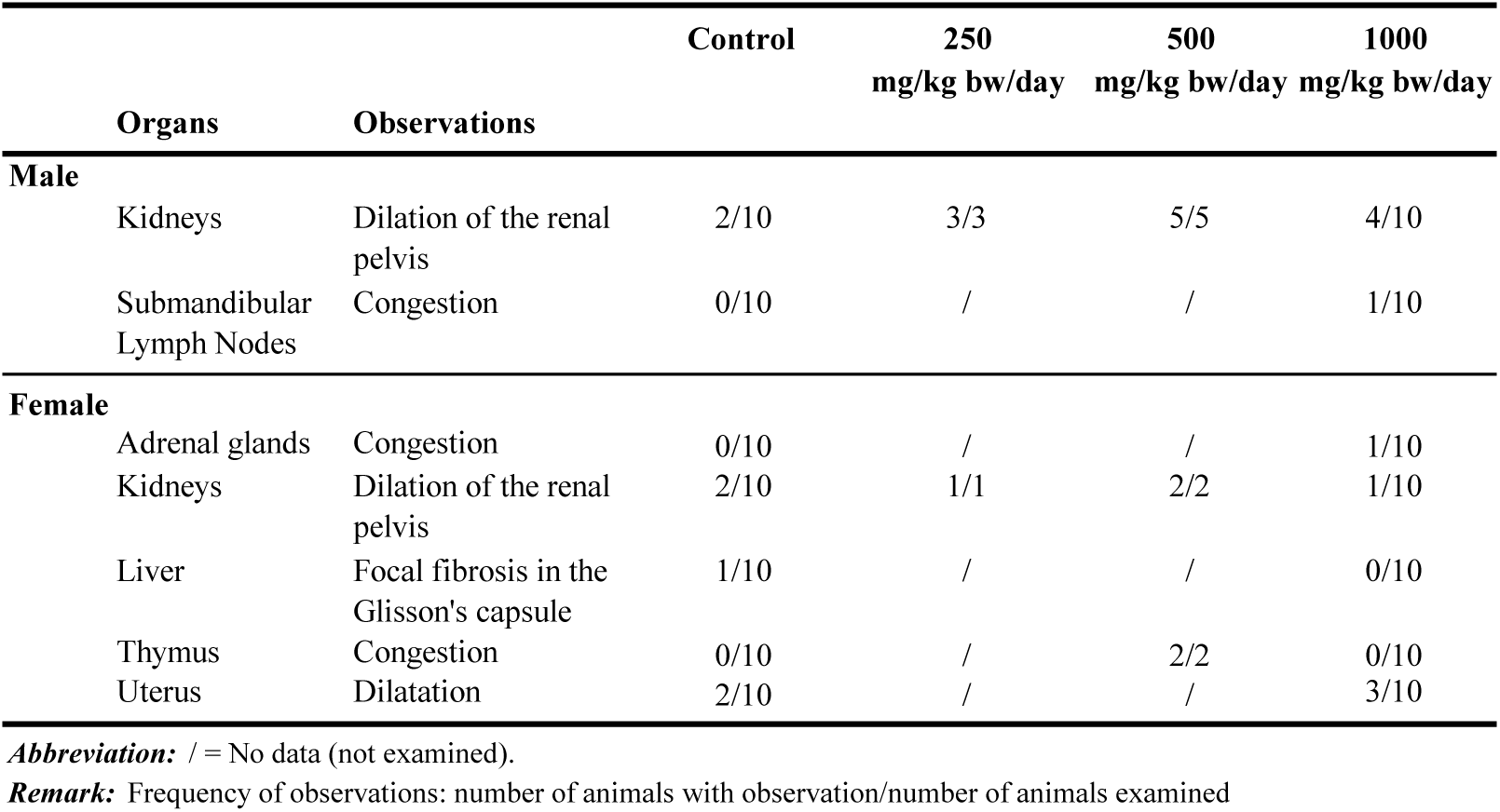
*Anaerostipes caccae* CLB101 90-day Histopathology Findings Summary.

## 4. Discussion

The in vivo comet assay results were clearly negative in the stomach and duodenal tissues, which had direct exposure to the test item, based on percent tail DNA values. Regarding the liver findings, while the lack of statistically significant results and dose-related increases in percent tail DNA suggest a lack of genotoxicity, they do not allow for the interpretation that direct or indirect liver exposure was achieved. Therefore, due to the lack of evidence of liver exposure, the comet assay cannot be considered clearly negative concerning the liver. At the same time, since the limit dose was applied in this study, we consider these comet assay results to be acceptably negative, despite the lack of liver exposure evidence. The negative ghost cell and Trypan blue findings further indicate that no cytotoxicity took place at any dose level in any tissue. The lack of genotoxicity found in mice in the micronucleus study, when bone marrow exposure was demonstrated by the slight, but statistically significant, reduction in PCE to total erythrocyte ratio in the high-dose group, supports the lack of genotoxicity seen in rats in the comet assay. Despite the possible interspecies differences and the lack of toxicokinetic understanding regarding the test item and/or its metabolites in each species, we considered that the combined results adequately demonstrate an overall lack of genotoxic potential of the test item.

The 90-day study did not reveal any adverse effects in the daily and weekly clinical observations or FOB. There were multiple statistically significant but minor hematology and blood coagulation changes relative to the control. Males in the high-dose group had lowered mean lymphocyte counts, and females in the mid-dose group had the same, in addition to lowered mean lymphocyte percentage. High-dose males also had lower mean platelet count and a slightly shorter mean activated partial thromboplastin time. Low- and mid-dose females had higher mean percentages of neutrophil granulocytes. All female test groups had higher mean corpuscular volume (MCV) and lower mean corpuscular hemoglobin concentration. Mid-dose females alone had higher hematocrit, while high-dose females alone had a lower mean percentage of monocytes. Despite the statistical significance, all the individual hematology and blood coagulation values remained well within or very close to the historical control ranges, and there were no correlated clinical chemistry or organ pathology changes. Therefore, these changes were considered minor and of no toxicological relevance. While the MCV values were nearly equal among all groups, the highest value was measured in the high-dose group. As such, it is unclear if there was a dose-dependent increase in MCV. Regardless, in no case were there accompanying macroscopic or histological findings, specifically hepatic, renal, or pancreatic, which might be expected if the aforementioned clinical pathology findings were toxicologically significant. Therefore, they were considered to have little or no toxicological relevance.

Clinical chemistry analyses also revealed statistically significant findings in male and female animals relative to the controls. In males, the low-dose group’s mean alanine aminotransferase (ALT) was elevated, the mid-dose group had a low mean concentration of glucose, the high-dose males had a lower mean potassium concentration, and the low and mid-dose groups had lower calcium compared to controls. In low and high-dose females, the mean concentration of inorganic phosphorus (Pi) was lowered, and high-dose females had elevated mean activity of aspartate aminotransferase. In all the female test groups, the mean concentration of chloride was increased, but it was not dose-dependent. All of these clinical chemistry findings were well within the historical control ranges. Therefore, they were considered toxicologically and biologically irrelevant. The serum concentration of thyroid-stimulating hormone, free triiodothyronine (FT3), and free thyroxine (FT4) of all test groups were comparable to the controls, except males in the mid and high-dose groups had increased FT3, while males in the high-dose group had high FT4. As there were no correlative macroscopic or histological findings in the pituitary, midbrain sections containing the hypothalamic region, or thyroid, the effects were considered sporadic occurrences and not toxicologically significant.

The lesions observed during the gross pathological examinations occurred in all groups, test and control, without dose responses and without histopathological corollaries. Based on the laboratory’s experience and historical control data, these lesions were consistent with common species-specific, developmental, or neuro-hormonal disorders in untreated experimental rats (of this strain and age). The two findings of thymus congestion in the mid-dose female group—the only gross findings not also found in the control and/or high-dose groups—were, upon histopathological exam, considered to have been caused by a circulatory disturbance developed during euthanasia and exsanguination. Therefore, they were also considered toxicologically irrelevant and unrelated to test item administration. The mean organ weight differences compared to controls were exclusive to females and included a number of statistically significant but sporadic changes across the dose groups that remained well within historical control ranges, were not dose-related, and/or were not correlated with other study findings. Additionally, all mid- and high-dose female thymus weights, absolute and relative to both the body and brain weights, decreased statistically significantly and dose-dependently. One high-dose thymus weight (relative to the body weight) fell below the historical control data, but there were no consistent correlative histopathology or clinical chemistry findings. Thus, none of the organ weight findings were considered to be test item-related or toxicologically relevant.

Very little supportive safety data regarding related species were identified, lending importance to this novel genotoxicity and toxicity assessment of *A. caccae* CLB101, specifically, and of clostridium cluster XIVa butyrate-producing colonic bacteria, generally. That said, another early safety assessment by Zhang et al. (2022) on *Rosburia intestinalis*, a related, butyrate-producing clostridium cluster XIVa bacteria, generally reflects the lack of toxicity findings identified in the current work for *A. caccae*. Their OECD 407 28-day repeated-dose oral toxicity study determined a NOAEL of 1.32 × 10^9^ CFU/kg/day, noting that they did not clarify which dose group determined the NOAEL, or for that matter, explicitly define the dose groups (they were defined as relative to the empirical dose of oral probiotics in humans (10^8^–10^9^ CFU or 1.43 × 10^6^–1.43 × 10^7^ CFU/kg/day in a 70 kg individual): control, 0; low-dose, 66–657 times; mid-dose, 77–769 times; and high-dose, 92– 923 times). No adverse effects were reported at any dose. A reduction of uric acid (UA) was noted in all dose groups, which was discussed as statistically significant although it is unclear if all or just the two higher dose groups were considered statistically significant. These results correlated with their OECD 423 acute oral toxicity study in mice which also had no adverse effects except a significantly reduced UA and an in vitro study (no data provided) they discussed, which indicated that *R. intestinalis* can degrade UA in the surrounding environment by 60% in 24 hours. The authors state that they are unsure of the mechanism of UA degradation (Zhang et al., 2022). The current 90-day study on *A. caccae* CLB101 did not measure UA levels, but interestingly, as discussed earlier, He et al. (2022) also found reduced UA levels when they fed inulin-type prebiotics to patients with renal failure and postulated that *A. caccae* may have had an indirect role in this reduction via cross-feeding with urate-reducing bacteria and by modulating the function of the intestinal urate exporter ABCG2 to potentiate extrarenal UA excretion (He, 2022). Further studies are required to confirm *A. caccae’s* ability to reduce UA in rats and to uncover the mechanism(s) of action.

### 4.1 Conclusions

Under the conditions of the experiments described in these studies, no genotoxicity was observed in the in vivo comet assay or in the in vivo micronucleus assay. Moreover, no findings of toxicological relevance were found in the 90-day repeated dose oral toxicity study in Han:WIST rats, in which a NOAEL of 1000 mg/kg bw/d for both sexes (equivalent to 1.9 x 1011 CFU/kg bw/d), the highest dose tested, was determined. All studies used Mitsuoka buffer, in which cell viability has been previously determined to be ∼32% after one hour. Thus, the equivalent NOAEL for living cells would be ∼6.1 x 1010 CFU/kg bw/d. The absence of signs of toxicity presented in these studies indicates a promising potential for the use of A. caccae CLB101 in food.

## Supporting information

Supplemental Table 1 Food Consumption

Supplemental Table 2 Body Weight Gain

## Acknowledgements

The authors thank the following individuals for their contributions to the work: Míra Andorka, Csaba Berczelly, Cecília Szijártó Bándiné, Ibolya Bogdán, Katalin Csendes, Tímea Csörge, Erika Dobos, Zsuzsanna Frank, Zoltán Gaál, Irén Somogyi Háriné, Ildikó Hermann, Brigitta Horváth, Istvánné Horváth, Zsolt Hummel, Petra Jáger-Gaál, Ágota Jó, Bálint Zsolt Juhari, Anita Klucsik, Dávid Kovács, Klára Fritz Kovácsné, Mónika Kozma, Nóra Pongrácz Kurdiné, Bence Küronya, Adrienn Laczó, Anikó Légrádi-Maurer, Marcell Madár, Máté Madár, Viktória Matina, Edit Kővári Mesterháziné, Timea Molnár, Dániel Németh, Barbara Palombi, Anikó Renkó, Fruzsina Ritter-Tóth, Szilvia Simai, Sebestyén Simon, Zsuzsanna Szabó, Dávid Szabó. Dávid Szabó, Mariann Lennert Szabóné, Mónika Oláh Szabóné, Olga Szász, Márta Tenk, Erika Misku Vargáné, Levente Zoltán.

## Statements and Declarations

### Ethical Considerations

The animal studies were approved by the Institutional Animal Care and Use Committee of Toxi-Coop Zrt. The 90-day study was additionally conducted according to the National Research Council Guide for Care and Use of Laboratory Animals (National Research Council, 2011) and in compliance with the principles of the Hungarian Act 2011 CLVIII (modification of Hungarian Act 1998 XXVIII) and Government Decree 40/2013 regulating animal protection.

### Declaration of Conflicting Interest

Authors Vickie Modica and Amy Clewell are salaried employees of AIBMR Life Sciences, Inc. (Seattle, WA, USA). AIBMR was contracted by the study sponsor as an independent third party to determine appropriate study protocols and dose selections, place the studies, approve the study plans, to monitor the toxicological studies herein described, to analyze and interpret the resulting data, and to prepare the manuscript. Author John Endres is a former salaried employee of AIBMR Life Sciences, Inc.

Author Gábor Hirka is owner and Managing Director at Toxi-Coop Zrt. (with test facilities in Budapest and Balatonfüred, Hungary); authors Adél Vértesi, Erzsébet Béres, and Ilona Pasics Szakonyiné, are salaried employees of Toxi-Coop; and author Róbert Glávits is an independent contractor to Toxi-Coop. Toxi-Coop was contracted by AIBMR to develop the study plans and conduct, analyze, and interpret and report the results of the toxicological studies herein described.

The authors declare no additional conflicts of interest in regard to the research, authorship, and/or publication of this article.

### Funding Statement

The authors disclose that financial support for the research described herein was provided by ClostraBio, Inc. Chicago, IL.

